# *In Vitro* Neutrophil-Bacteria Assay in Whole Blood Microenvironments with Single-Cell Confinement

**DOI:** 10.1101/2024.01.22.576723

**Authors:** Chao Li, Nathan W. Hendrikse, Zach Argall-Knapp, Makenna Mai, Jun Sung Kim

## Abstract

Blood is a common medium through which invasive bacterial infections disseminate in the human body. *In vitro* neutrophil-bacteria assays allow flexible mechanistic studies and screening of interventional strategies. In standard neutrophil-bacteria assays, both the immune cells and microorganisms are typically interrogated in an exogenous, homogeneous, bulk fluid environment (e.g., culture media or bacterial broth in microtiter plates), lacking the relevant physicochemical factors in the heterogenous blood-tissue microenvironment (e.g., capillary bed) with single-cell confinement. Here we present an *in vitro* neutrophil-bacteria assay by leveraging an open microfluidic model known as “μ-Blood” that supports sub-microliter liquid microchannels with single-cell confinement. In this study we compare the exogenous and endogenous fluids including neutrophils in RPMI (standard suspension cell culture media) and whole blood in response to *Staphylococcus aureus* (*S. aureus*, a gram-positive, non-motile bacterium) in phosphate buffered saline (PBS), Mueller Hinton Broth (MHB), and human serum. Our results reveal a significant disparity between the exogenous and endogenous fluid microenvironments in the growth kinetics of bacteria, the spontaneous generation of capillary (i.e., Marangoni) flow, and the outcome of neutrophil intervention on the spreading bacteria.

## 1. Introduction

As the first line of defense in the innate immune system, neutrophils swiftly respond and migrate toward the infection niche and intervene with multiple anti-infective strategies^[1,2]^ including phagocytosis (i.e., internalization and killing of pathogens),^[3]^ degranulation (i.e., release of antimicrobial granules),^[4]^ and the formation and release of neutrophil extracellular traps (NETs) - a web-like structure composed of chromatin, histones, and antimicrobials.^[5]^ Studies of neutrophil-pathogen interactions provide insights into new interventional strategies and treatment targets that may better prevent and cure infections.

In the war between neutrophils and invading pathogens, the first event in neutrophil intervention is the recruitment of neutrophils to the infection niche via migration.^[6,7]^ In light of the importance of this initial event, different neutrophil migration mechanisms [e.g., chemotaxis,^[8–10]^ swarming^[11–15]^] have been studied with both *in vivo* [e.g., mouse,^[16]^ zebrafish^[17]^] and *in vitro* models.^[18–20]^ While *in vivo* animal models provide a complete system, they are limited by adoption/implementation barriers, cost, throughput, and most importantly are non-human, limiting their relevance to human disease. In comparison, *in vitro* models, especially microfluidics-based systems, provide a venue to overcome some of these limitations. Importantly, *in vitro* models can utilize human primary samples (e.g., blood, tissue, microbial isolates directly from patients) to bridge the gap between *in vitro* and *in vivo* to provide a promising pipeline toward personalized/precision medicine.^[21]^

To date, *in vitro* immune-pathogen studies are typically performed in an artificial culture media at bulk scale (i.e., a space much larger than single cell or bacterium), exposing both immune cells and microorganisms to an exogenous fluid microenvironment that lacks many of the relevant physicochemical factors present *in vivo*. The altered microenvironment can result in a significant loss of *in vivo* mechanisms (and thus information) and inconsistencies of the assays from different sources, e.g., experiment to experiment, operator to operator, and lab to lab.^[22]^ Recently, we developed an *in vitro* neutrophil functional assay known as “μ-Blood”.^[23]^ μ-Blood allows interrogation of primary neutrophils directly in a microliter of unprocessed whole blood with single-cell confinement, better preserving the *in vivo* donor-specific information with improved assay consistency.

In this study, we investigate neutrophil-bacteria (*S. aureus*) interactions in μ-Blood by comparing the standard assay fluid microenvironment with the whole blood microenvironment for their influence on both neutrophils and bacteria. Neutrophils are either kept in whole blood or isolated from whole blood and then resuspended in RPMI [i.e., Roswell Park Memorial Institute (RPMI) 1640 Medium. Bacteria (*S. aureus*) are cultured in MHB (i.e., Mueller Hinton Broth) and resuspended in PBS, MHB, and human serum. The μ-Blood assay results reveal distinctive outcomes of neutrophil intervention on the invading bacteria in the single-cell confined space, depending on the growth kinetics of bacteria at the infection niche and the spontaneous generation of Marangoni flow - a capillary convection driven by a surface/interfacial tension differential in a fluid. In a quasi-stationary fluid microenvironment with mild to no Marangoni flow, neutrophils show successful intervention in response to the invading bacteria in the confined space - physically pushing the bacteria out and back to the infection niche, phagocytizing bacteria in their way, killing the internalized bacteria, and releasing granules and NETs. By contrast, getting carried in Marangoni flow triggered by the growth of bacteria at the infection niche, neutrophil intervention gets impaired and reversed with the responding neutrophils getting physically pushed back and the bacteria getting directly pumped into the immune niche. In addition, *S. aureus* show different growth kinetics cultured in MHB and human serum, which determines the generation and intensity of Marangoni flow during neutrophil intervention through the confined space. These results highlight the necessity to investigate immune-pathogen interactions *in vitro* in a microenvironment that recapitulates the highly heterogeneous components, structures, and fluid dynamics as observed *in vivo*.

## 2. Results and Discussion

### 2.1. The Confined Whole Blood Microenvironment in μ-Blood

The design philosophy of μ-Blood^[23]^ (Figure 1A) is to establish an immune cell functional assay with an autologous whole blood (i.e., an endogenous fluid) microenvironment as opposed to an artificial culture media (i.e., an exogenous fluid) microenvironment primarily adopted in the standard cell culture/assays.

**Figure 1.**
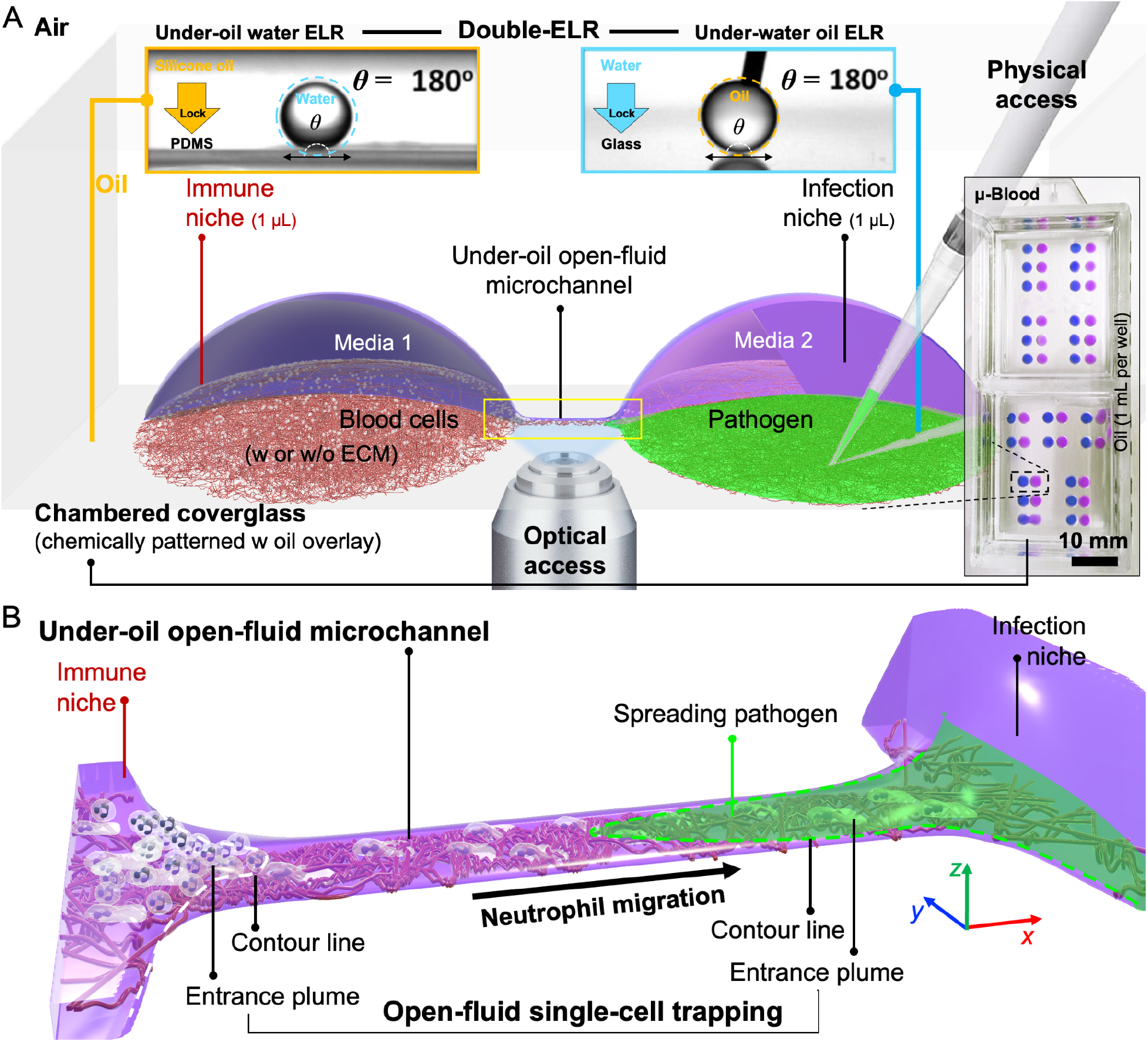
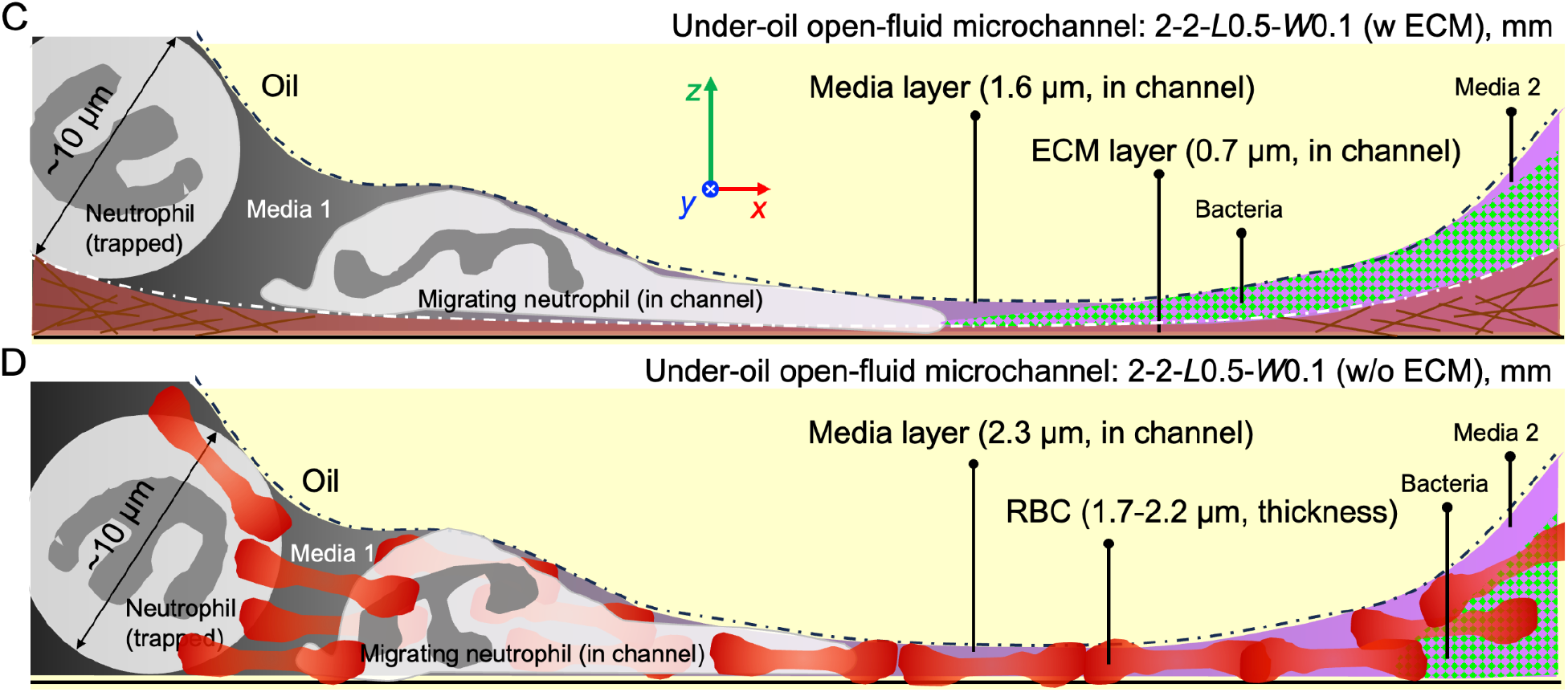
The under-oil open-fluid microchannel in μ-Blood with single-cell confinement. (A) A 3D model of the under-oil open-fluid microchannel 2-2-*L*0.5-*W*0.1 (w ECM). The coverglass surface is chemically patterned - background [a monolayer of polydimethylsiloxane (PDMS)-silane covalently bonded to the surface); patterned areas [oxygen (O_2_) plasma-treated)] (Movie S1). The surface is overlaid with silicone oil (5 cSt, 1.6 mm in thickness for 1 mL per well). (Insets) Contact angle images of under-oil water ELR (on PDMS-grafted areas) and under-water oil ELR (on O_2_ plasma-treated areas), together for Double-ELR. Droplets are 3 μL in volume. The microchannel is highlighted by the yellow solid line box. (B) Zoomed-in image of the microchannel. The entrance plumes are defined by the contour line (colored dashed line) at the immune niche and the infection niche. (C) Side view (xz plane) of the microchannel with ECM coating showing the thickness of the ECM (0.7 μm) and media layer (1.6 μm). Neutrophils (∼10 μm in diameter when stand-by) get trapped at the entrance plume. *S. aureus* (0.5 to 1.5 μm in diameter) spreads into the microchannel from inoculation. Neutrophils migrate into the microchannel, taking a flattened and elongated cell shape. (D) Side view of the microchannel without ECM coating showing the thickness of the media layer (2.3 μm). RBCs (7.5 to 8.7 μm in diameter and 1.7 to 2.2 μm in thickness) spread into the microchannel from whole blood loading, forming a monolayer with the RBC’s disc in parallel to the substrate surface.

The technical basis on which μ-Blood is developed is an extreme wettability phenomenon known as Exclusive Liquid Repellency (ELR)^[24]^. ELR allows a liquid (e.g., aqueous media, plasma/serum, whole blood) to be inherently (i.e., surface-texture and surfactant independent) and absolutely repelled on a solid surface (with Young’s contact angle *θ* = 180°) when exposed to a secondary immiscible liquid phase (e.g., oil).^[24–26]^ Double-ELR is an expansion of ELR that integrates under-oil water ELR and under-water oil ELR on a chemically patterned surface (Figure 1A, insets).^[25]^ Double-ELR enables a new sub-branch in open microfluidics - ELR-empowered Under-oil Open Microfluidic Systems (UOMS).^[26]^ In ELR-empowered UOMS, micrometer-scale under-oil open-fluid microchannels can be prepared via a unique liquid handling technique known as under-oil sweep distribution (Figure S1, Movie S1, Experimental Section). To perform under-oil sweep distribution, a hanging drop of media (any size compared to the surface pattern) gets dragged across the Double-ELR surface by a pipet (Movie S1). A specific volume of the media [with a nearly constant height-to-width (*H*/*W*) 1/13 of the media layer]^[26]^ is spontaneously distributed on the patterned areas. When the hanging drop is outside of the patterned areas it is absolutely repelled, limiting sample loss and device fouling.

Defined by this Double-ELR physics and the unique three-dimensional (3D) geometry of the under-oil microchannels, open-fluid single-cell trapping (Figure 1B, C) can be achieved with a channel width (100 μm) much larger than the size of the target cells, e.g., white blood cells, 7 to 30 μm in diameter; RBCs, 7.5 to 8.7 μm in diameter and 1.7 to 2.2 μm in thickness; platelets, 3 to 4 μm in diameter and *S. aureus*, 0.5 to 1.5 μm in diameter. In this study, we used under-oil microchannels with the channel dimensions written in the format of 2-2-*L*0.5-*W*0.1 [w or w/o extracellular matrix (ECM)] for 2 mm in diameter of the immune niche and infection niche spots, 0.5 mm in channel length (*L*), and 0.1 mm in channel width (*W*). With a given spot size, channel length, and channel width, the channel height - i.e., the ECM layer thickness + the media layer thickness for the microchannels with ECM coating or the the media layer thickness for the microchannels without ECM coating - can be known from the calculation of Laplace pressure equilibrium.^[23]^

For 2-2-*L*0.5-*W*0.1 (w ECM), the ECM layer thickness is 0.7 μm with a media layer thickness of 1.6 μm on top (Figure 1C).^[23]^ The 2-2-*L*0.5-*W*0.1 (w ECM) microchannels can effectively trap blood cells on the immune niche spot at the entrance plume. *S. aureus* can spread into the microchannel to a varying extent depending on the inoculum density and volume on the infection niche spot. Neutrophils can only move into the microchannel via active cell deformation and migration with a noticeably flattened and elongated cell shape (Figure 1C).^[23]^ For 2-2-*L*0.5-*W*0.1 (w/o ECM), the media layer thickness is 0.7 μm + 1.6 μm = 2.3 μm, which leads to a monolayer of RBCs spreading into the microchannel from whole blood loading with the RBC’s disc in parallel to the substrate surface (Figure 1D). Due to the occupation of the microchannel space by RBCs, *S. aureus* gets excluded out of the microchannel and trapped on the infection niche spot. The open-fluid single-cell trapping function lays the foundation for running *in vitro* immune-pathogen assays with a confined whole blood microenvironment and various pathogens.

### 2.2. The Key Neutrophil-Pathogen Control Events in Response to Living Bacteria in Under-Oil Open-Fluid Microchannels

The primary function of neutrophils is the immediate response to and neutralization of invasive microorganisms (e.g., fungi, bacteria, or viruses). In this experiment, we demonstrate neutrophil recruitment and the neutrophil-pathogen control events in response to living bacteria [green fluorescent protein (GFP)-labeled *S. aureus*] in the under-oil open-fluid microchannels with single-cell confinement (Figure 2, Movie S2). We first put isolated neutrophils (Experimental Section) in standard RPMI media (1 × 10^4^ cells/μL) in response to GFP-labeled *S. aureus* (1 × 10^6^ CFU/μL) in PBS using the microchannels 2-2-*L*0.5-*W*0.1 (w ECM) with single-oil (silicone oil, 5 cSt) overlay (Figure 2A).

**Figure 2.**
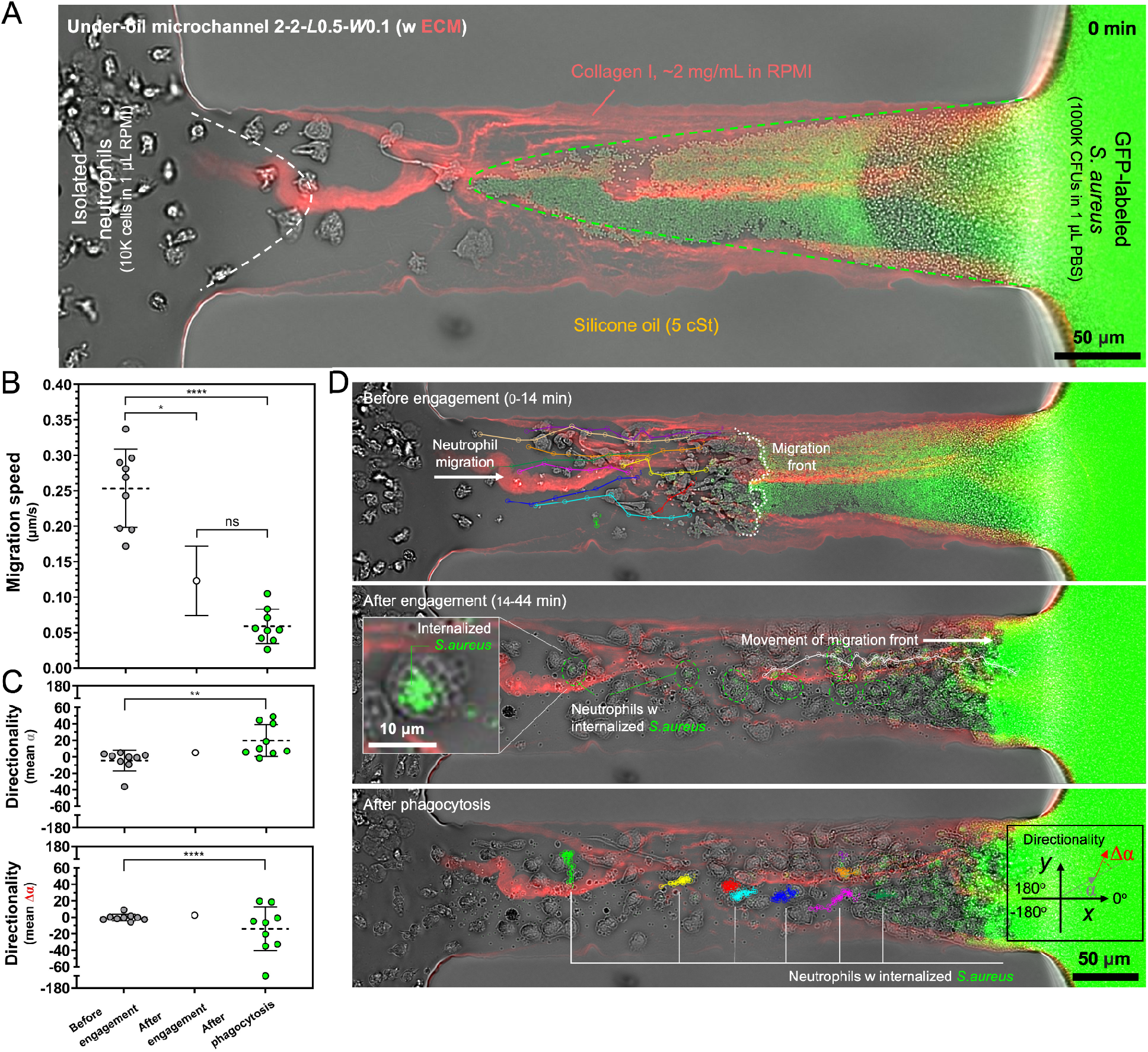
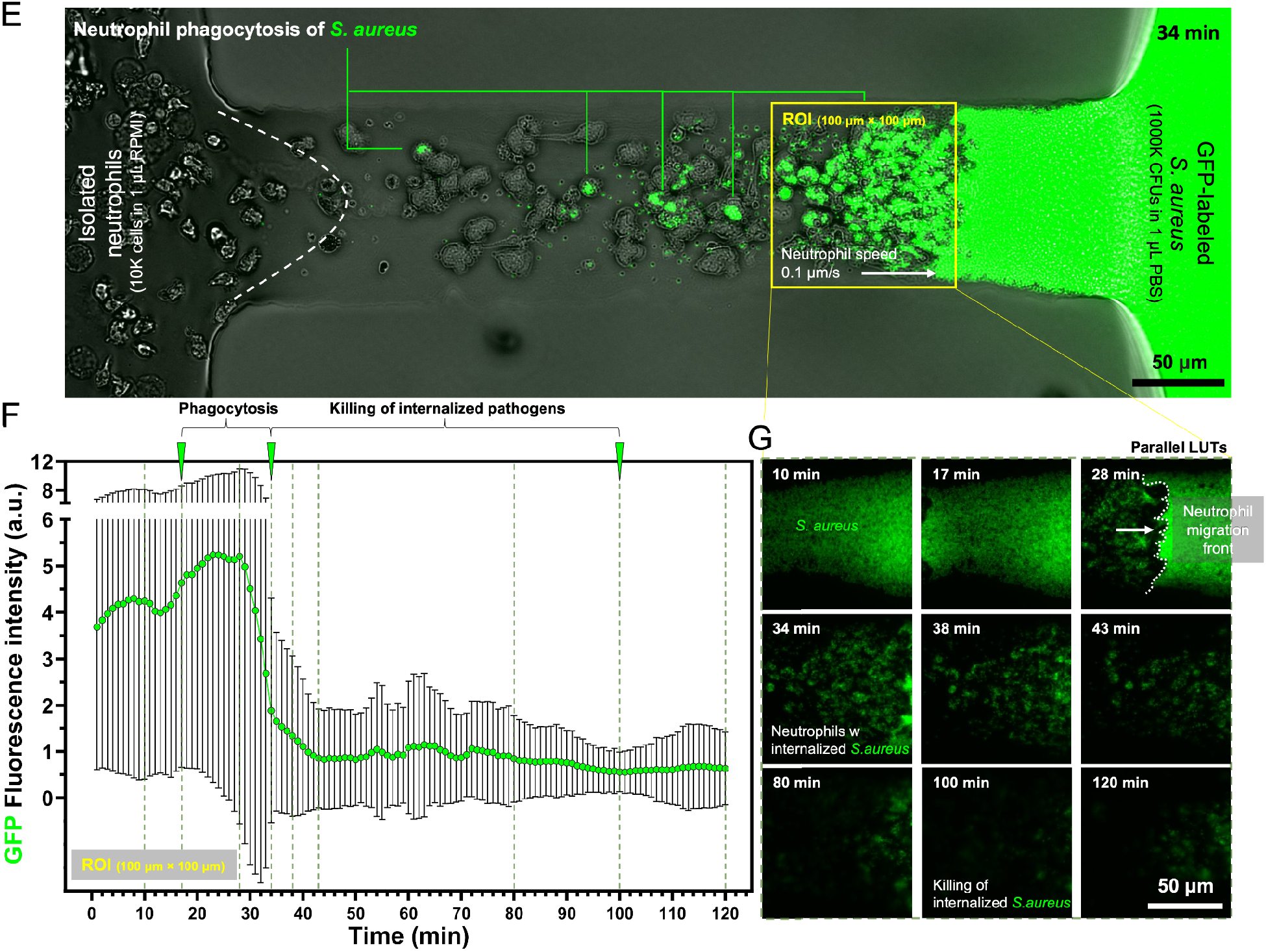
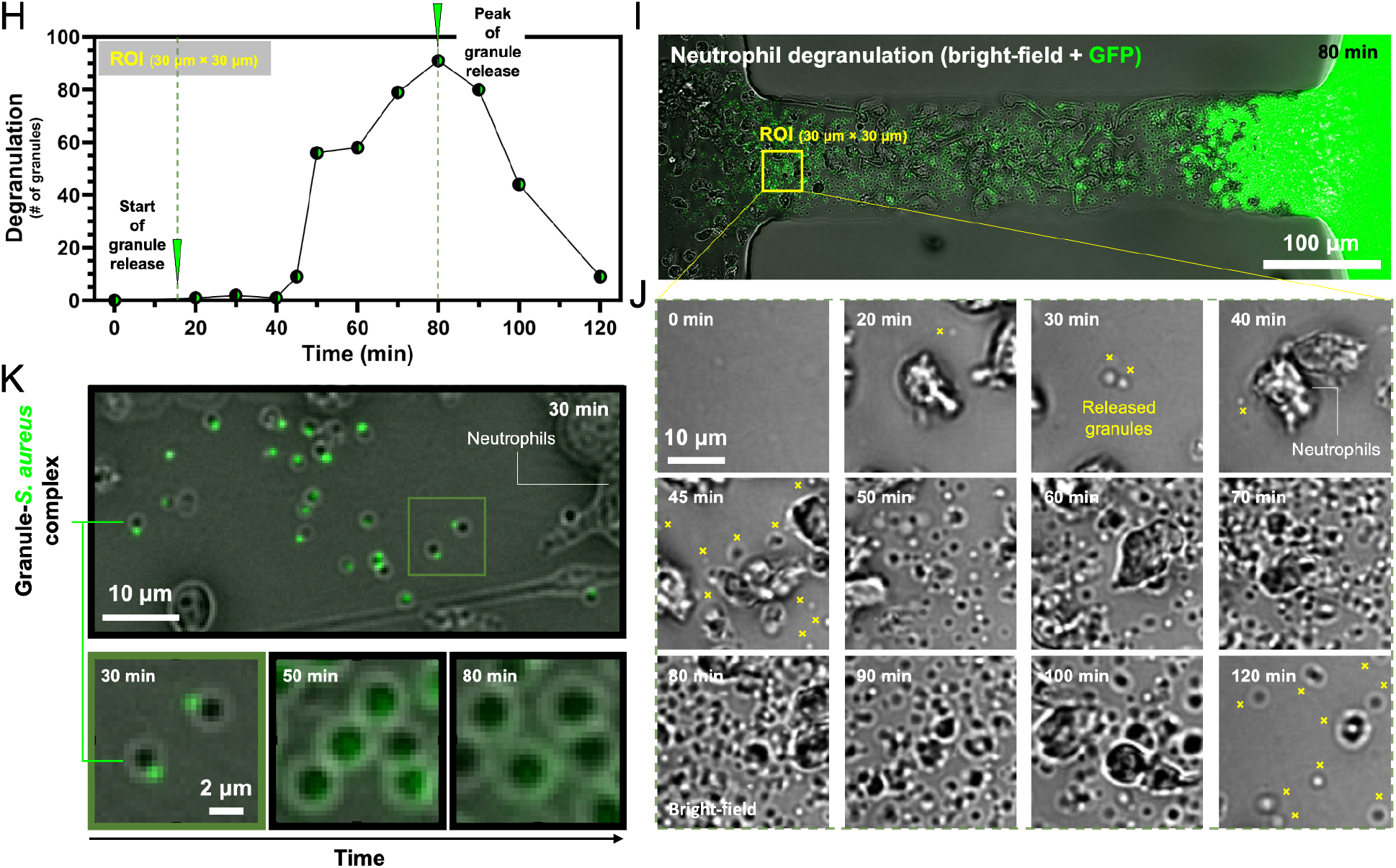
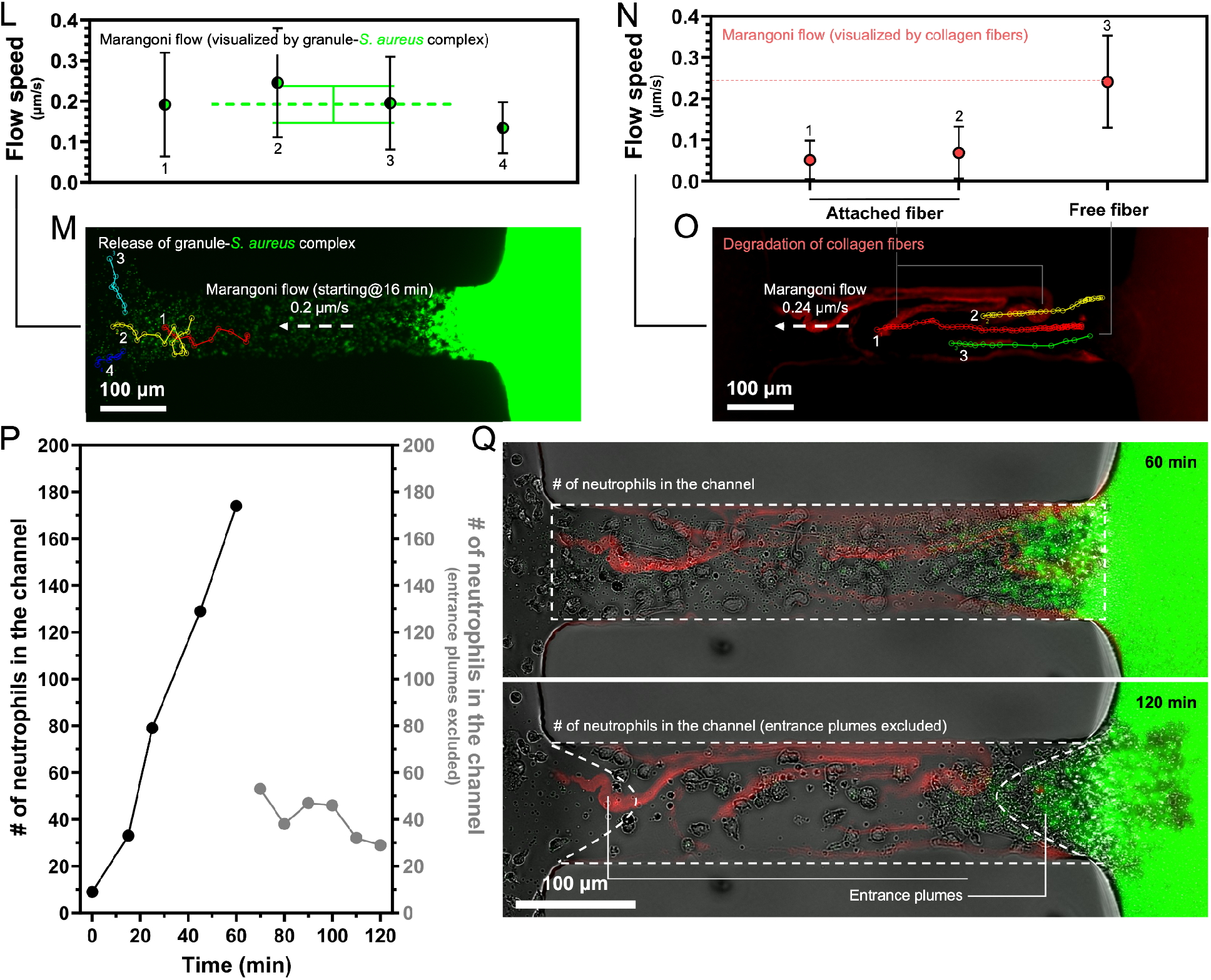
Isolated neutrophils respond to *S. aureus* (GFP-labeled) in an under-oil open-fluid microchannel. A series of pathogen-control events of neutrophils in 2 h were captured and quantified. (A) A top view (xy plane) of the under-oil microchannel [2-2-*L*0.5-*W*0.1 (w ECM - collagen I, ∼2 mg/mL in RPMI)] with the immune niche (neutrophils in 1 μL RPMI, 10K cells/μL) on the left and the infection niche (*S. aureus* in 1 μL PBS, 1000K CFU/1 μL) on the right. The dashed lines highlight the entrance plumes at 0 min (i.e., right after sample loading). (B) Neutrophil migration speed and (C) migration directionality (α - the angle between the current displacement vector and the x direction; Δα - the angle between the current and the previous displacement vectors) analyses before/after encounter with the bacteria and after phagocytosis. (D) The representative microscopic images (composite of bright-field, GFP, and RFP) with cell tracks corresponding to (B) and (C). (Inset) Close-up of a neutrophil with internalized *S. aureus*. (E) A microscopic image (composite of bright-field and GFP) showing neutrophil phagocytosis of *S. aureus* at 34 min. The yellow box shows the ROI (100 μm × 100 μm) of the phagocytosis and killing analyses. (F) Change of the mean GFP fluorescense intensity in the ROI in (E). (G) The representative microscopic images with parallel lookup tables (LUTs) for a comparable intensity. (H) Degradation analysis, i.e., the number of released granules as a function of time. (I) A microscopic image (composite of bright-field and GFP) showing the released granules at its peak intensity at 80 min. The yellow box shows the ROI (30 μm × 30 μm) of the degranulation images in (J). (J) The representative microscopic images (bright-field) showing the released granules - highlighted by the yellow cross. (K) Close-ups of the granule-*S. aureus* complex with an asymmetrical structure and the evolution over time. (L) and (M) A backflow (with its direction from the infection niche to the immune niche) was detected at 16 min for 0.2 μm/s, visualized by the movement of the fluorescent granule-*S. aureus* complex. (N) and (O) Degradation of ECM fibers and the backflow visualized by the movement of the ECM fibers for 0.24 μm/s (free fiber). (P) The number of neutrophils in the channel (left) and the entrance plumes excluded (right) over time. (Q) The representative microscopic images showing the reduced neutrophil count in the channel after 60 min due to the influence of the backflow. Error bars are mean ± S.D. from ≥3 replicates. \**P* ≤ 0.05, \*\**P* ≤ 0.01, \*\*\**P* ≤ 0.001, and \*\*\*\**P* ≤ 0.0001. “ns” represents “not significant”.

In the first 2-h window after sample loading, a full set of neutrophil-pathogen control events were captured. Right after sample loading, neutrophils responded to and migrated toward the bacteria in the microchannels at an average speed of 0.25 μm/s on ECM before the cells encountered the bacteria (Figure 2B-D). At 14 min, the migration front of neutrophils engaged the bacterial entrance plume in the microchannel. After the encounter, the neutrophil migration speed dropped to 0.12 μm/s due to the physical resistance from the bacterial layer in the microchannel (Figure 1C). Neutrophils kept migrating forward, pushing bacteria backward and out of the confined space, in the meanwhile, phagocytizing the bacteria in their way. Right after phagocytosis, neutrophils stopped migrating, losing their speed and directionality (Figure 2B, C, Movie S2). The apparent mutual exclusion between neutrophil migration and phagocytosis could be attributed to the competing consumption of the intracellular messenger - phosphatidylinositol (4,5)-bisphosphate (PIP2) - for migration/polarization PIP2 → PIP3 [phosphatidylinositol (3,4,5)-trisphosphate];^[8,9]^ for phagocytosis PIP2 → IP3 (inositol 1,4,5-trisphosphate) + DAG (diacylglycerol).^[27]^

After phagocytosis, the internalized bacteria started losing their GFP fluorescence over time, which indicates neutralization and killing of the bacteria inside neutrophils (Figure 2 E-G).^[3]^ In a region of interest (ROI) next to the infection niche, the killing of internalized bacteria took 1 h. During almost the same time window from 17 min to 120 min, degranulation of neutrophils occurred (Figure 2 H-K).^[28]^ During degranulation, neutrophil granules (2 μm in diameter as visualized under 20× magnification bright-field) were released, peaking at 60 min after the start of degranulation. Interestingly, the released neutrophil granules all formed an asymmetrical granule-*S. aureus* complex with 1 bacterial cell tethered to 1 granule (Figure 2K). Over time in 50 min, the tethered bacterium was apparently absorbed and neutralized by the granule, leading to reduced and homogenized GFP fluorescence and increased size of the neutrophil granule from 2 μm to 4 μm. During the phagocytosis and degranulation process, neutrophils got recruited continuously into the microchannel, showing little generation and release of NETs and channel clogging. In the following experiment (not in this experiment) with a DNA stain (Hoechst), NETs release can be identified after neutrophils migrated into the infection niche where neutrophils got exposed to bacteria in a significantly higher density. The distinct NET generation and release between neutrophils in the microchannel (little to none) and at the infection niche (significantly activated) indicates that NETosis is delicately modulated by multiple environmental factors^[29,30]^ including signaling cytokines,^[31]^ mechanical confinement^[32]^ and shear stress,^[33]^ and pathogen size^[34]^/density.^[35]^

Besides the canonical neutrophil-pathogen control events described above (i.e., neutrophil recruitment, phagocytosis, degranulation, and NETosis), an unexpected event - a “backflow” that is opposite to the direction of neutrophil migration, pointing from the infection niche to the immune niche - was captured during degranulation (Figure 2L, M). Visualized by the GFP fluorescence of the granule-*S. aureus* complex, the speed of the backflow was tracked to be 0.2 μm/s during the degranulation time window. Getting carried in the flow, the granule-*S. aureus* complex particulates got pumped directly into the immune niche (Movie S2). At 40 min after sample loading, neutrophils migrated through the microchannel and reached the infection niche (Figure 2D). At this time, the ECM fibers got snapped quickly in about 30 min at the connection between the microchannel and the infection niche (Movie S2), getting pushed into the microchannel (Figure 2N, O) at a speed of 0.24 μm/s (visualized by the motion of the free fibers) due to the backflow generated during the degranulation process. The apparent degradation and cutting of ECM are attributed to the release of ECM-remodeling enzymes (e.g., elastase)^[36]^ during NETosis at the infection niche. Influenced by the backflow, we observed a reduced neutrophil count in the microchannel at 60 min (Figure 2P, Q, Movie S2).

These neutrophil-pathogen control events and neutrophil-ECM interaction results revealed in this section showed that μ-Blood provides a quantitative and precisely controlled immune-pathogen assay microenvironment *in vitro*, recapitulating the highly heterogeneous components, structures, and fluid dynamics as seen *in vivo*. Compared to the homogenous, bulk fluid environment typically adopted in standard assays, the μ-Blood assay preserves and extracts more information relevant to *in vivo*.

### 2.3. Marangoni Flow in Exogenous Fluid Microenvironments - Isolated Neutrophils in RPMI versus *S. aureus* in PBS

Here a question is what is the backflow? In such a microfluidic channel without active pumping (i.e., liquid getting pumped into the channel via a fluid pump) there could be only two possible passive pumping mechanisms that generate a lateral flow (Figure S2). One is driven by a Laplace pressure differential with the flow direction pointing from high Laplace pressure (i.e., high curvature and/or interfacial tension) to low Laplace pressure (i.e., low curvature and/or interfacial tension). The other is driven by an interfacial tension differential known as Marangoni flow with the flow direction pointing from low interfacial tension to high interfacial tension. The microchannels in this experiment (2-2-*L*0.5-*W*0.1) were swept with a collagen solution (∼2 mg/mL in RPMI) first (Movie S1), leading to an ECM layer with a *H*/*W* ratio of 1/13 on the spots and in the microchannel.^[26]^ The volume of ECM on the spots (assuming taking a spherical cap shape) is calculated to be 0.24 μL. After ECM sweeping and polymerization (1 h at room temperature), the spots were loaded with an extra 1 μL of volume for isolated neutrophils in RPMI and *S. aureus* in PBS under oil. The total volume (0.24 + 1 = 1.24 μL) leads to a spherical cap with a height of 0.69 mm and radius of curvature (*R*) of 1.07 mm. Due to the same loading volume (and thus curvature) and the close interfacial tension between silicone oil (5 cSt)-RPMI (*γ*_oil-RPMI_ = 39.9 mN/m) and silicone oil (5 cSt)-PBS (*γ*_oil-PBS_ = 40.2 mN/m) (Table S1), the two spots were well balanced after sample loading, which generates nearly no lateral flow from sample loading.

The experimental observation was consistent with the theoretical analysis above - the backflow was little to none right after sample loading. However, the backflow started to be detectable at 16 min, increasing over time up to 18 μm/s at 160 min (Movie S3). We can estimate the volume that gets pumped out of the infection niche in 160 min is <89 nL [i.e., 18 μm/s (the flow rate)^[26]^ × 515.2 μm^2^ (cross-section area of the microchannel) × 160 min = 89 nL]. Compared to the volume on the spot (1.24 μL), the volume loss (<89 nL) during the backflow is negligible, causing little change in the curvature of the sessile microdrop on the spot. In this circumstance, if we assume the backflow is Laplace-pressure driven, it requires an increase of the interfacial tension at the infection niche and/or a decrease of the interfacial tension at the immune niche over time. If we assume the backflow is Marangoni flow, the interfacial tension change is just opposite to the Laplace-driven flow, i.e., a decrease of the interfacial tension at the infection niche and/or an increase of the interfacial tension at the immune niche over time.

It has been reported that bacteria secrete a significant amount of biosurfactants during growth, which lowers the surface/interfacial tension of the culture fluid over time.^[37,38]^ This fact points the backflow mechanism to Marangoni flow. To test out this hypothesis, here we compare different inoculum densities of *S. aureus* at the infection niche and its influence on generation of the backflow (Figure 3). We tested three inoculum densities of *S. aureus* at 1000K, 100K, and 25K CFU/μL in 1 μL PBS. The immune niche remained the same for 10K isolated neutrophils in 1 μL RPMI.

**Figure 3.**
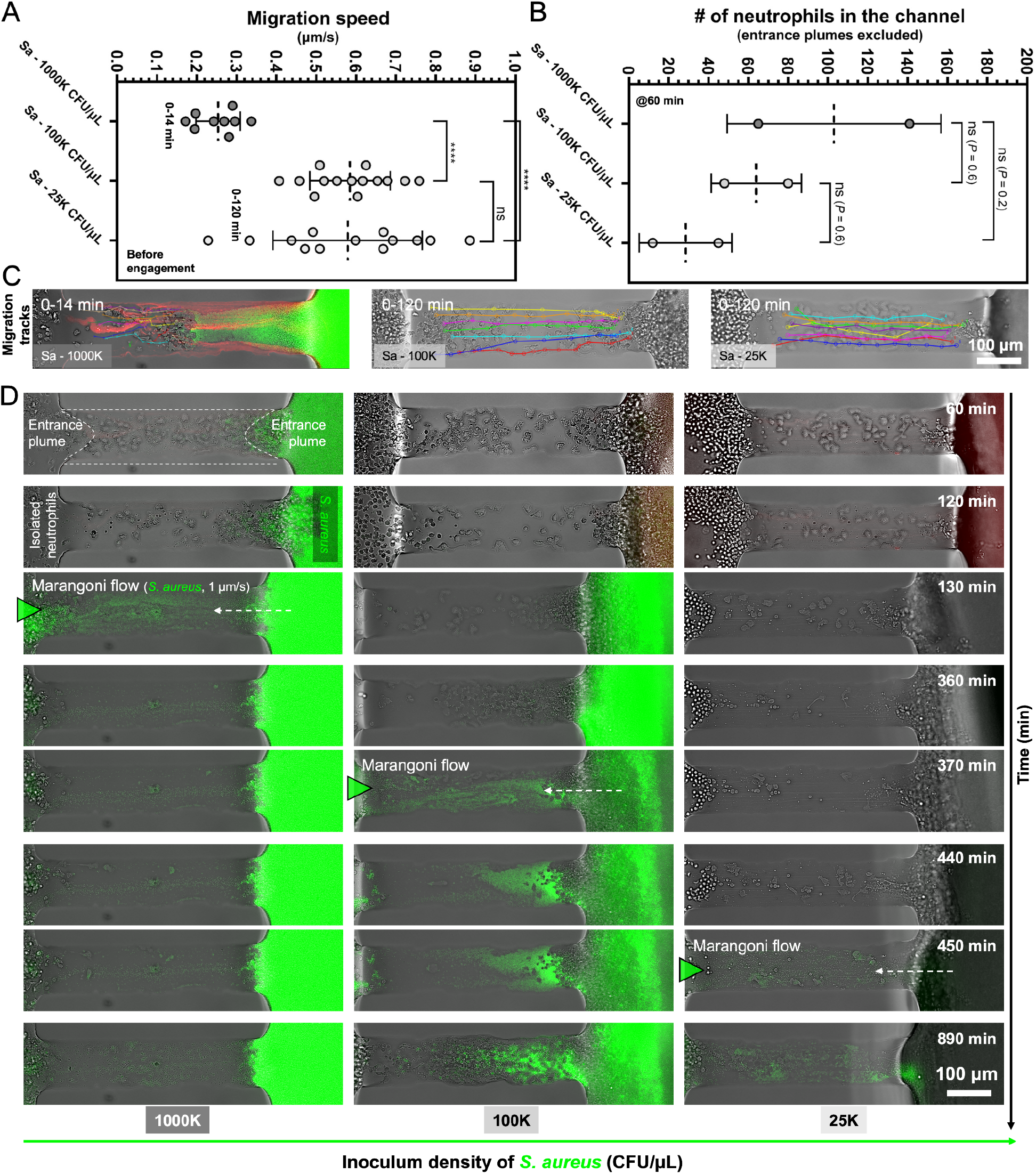
Influence of inoculum density of *S. aureus* on generation of backflow in the under-oil microchannel 2-2-*L*0.5-*W*0.1 (w ECM). (A) Neutrophil migration speed (before encounter with bacteria) and (B) the number of neutrophils in the channel (entrance plumes excluded) at 60 min in response to the three tested inoculum densities - 1000K CFU/μL, 100K CFU/μL, and 25K CFU/μL - in 1 μL PBS at the infection niche (the right spot). Neutrophils were seeded at the immune niche (the left spot) for 10K cells in 1 μL RPMI. (C) The representative microscopic images showing the cell tracks. With the inoculum densities lower than 1000K CFU/μL, no bacteria was found in the microchannel from sample loading. (D) Generation of the backflow (pointing from the infection niche to the immune niche) after 60 min. Low inoculum density led to a delayed generation of the backflow. The backflow was peaking at 130 min (1000K CFU/μL), 370 min (100K CFU/μL), and 450 min (25K CFU/μL), respectively, and visualized by the carryover streamlines of *S. aureus*. The burst of *S. aureus* carryover in the flow occurred fast from none in a time window of only 10 min. Error bars are mean ± S.D. from ≥3 replicates. \**P* ≤ 0.05, \*\**P* ≤ 0.01, \*\*\**P* ≤ 0.001, and \*\*\*\**P* ≤ 0.0001. “ns” represents “not significant”.

First, the results showed a distinct difference in neutrophil recruitment (Figure 3 A-C). High inoculum density of bacteria recruited more neutrophils but with reduced migration speed (0.25 μm/s) compared to the two lower inoculum densities (0.56 μm/s) (Movie S4). These results are consistent with the data from a parallel study of neutrophil migration in response to N-Formylmethionine-leucyl-phenylalanine (fMLP) - a bacterial peptide and standard for neutrophil chemotaxis study.^[39]^

As shown in the previous section, a backflow was visualized by the granule-*S. aureus* complex particulates starting at 16 min with a speed of 0.2 μm/s (Figure 2 L-O) and peaking at 130 min with a speed up to 1 to 18 μm/s (Figure 3D, Movie S3) from the 1000K CUF/μL inoculum density condition. The large range of the peak speed of the backflow is attributed to the varying levels of clutter density in the microchannel. An emptier channel after the neutrophils were purged and pushed out of the microchannel led to a higher flow speed due to the reduced hydrodynamic resistance. The two lower inoculum densities led to an obvious delay of the backflow (Figure 3D). The delayed generation of the backflow can be attributed to the increased time in bacterial growth and secretion of biosurfactants at the infection niche associated with a lower inoculum density.

### 2.4. Marangoni Flow in Exogenous Fluid Microenvironments - Isolated Neutrophils in RPMI versus Zymosan Particles in PBS

With the evidence from the inoculum density test in the section above, the backflow is clearly related to and influenced by the infection niche. Next we switched to dead pathogens - Zymosan particles (dead yeast cell walls so having no secretion of biosurfactants over time) to further confirm the mechanism of the backflow (Figure 4). In response to the dead yeast cell particles, no backflow was detected in 134 min after sample loading (Figure 4C, D, and F, Movie S5). The backflow was detected in the control group with isolated neutrophils in RPMI versus *S. aureus* in PBS for 0.62 μm/s at 48 min after sample loading (Figure 4C-E). This result rules out the change of interfacial tension over time (for 2 h) at the immune niche with isolated neutrophils in RPMI. Therefore, the generation of the backflow can be only caused by either an increased interfacial tension (if Laplace pressure-driven flow) or a decreased interfacial tension (if Marangoni flow) over time at the infection niche. Regarding the fact that bacteria secrete biosurfactants during growth, which reduces the interfacial tension at the infection niche over time,^[37,38]^ the backflow is identified as Marangoni flow caused and determined by the growth of bacteria at the infection niche.

**Figure 4.**
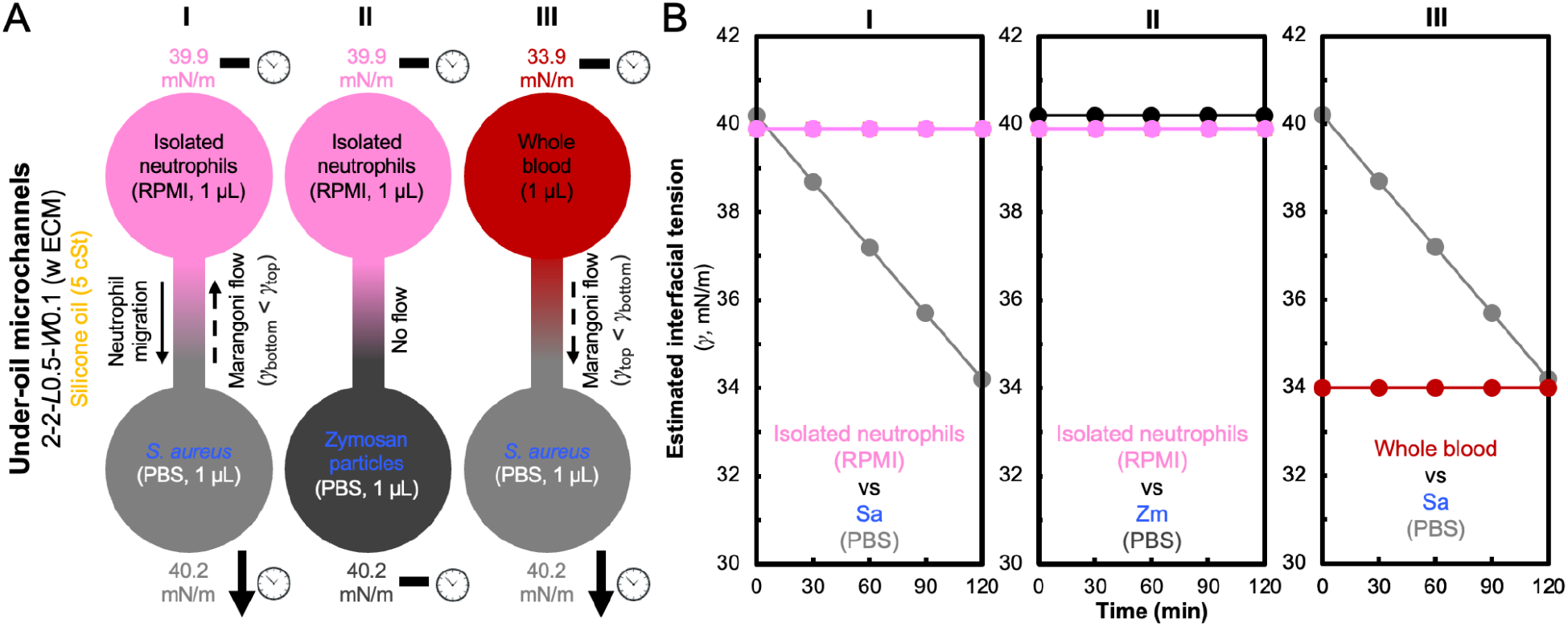
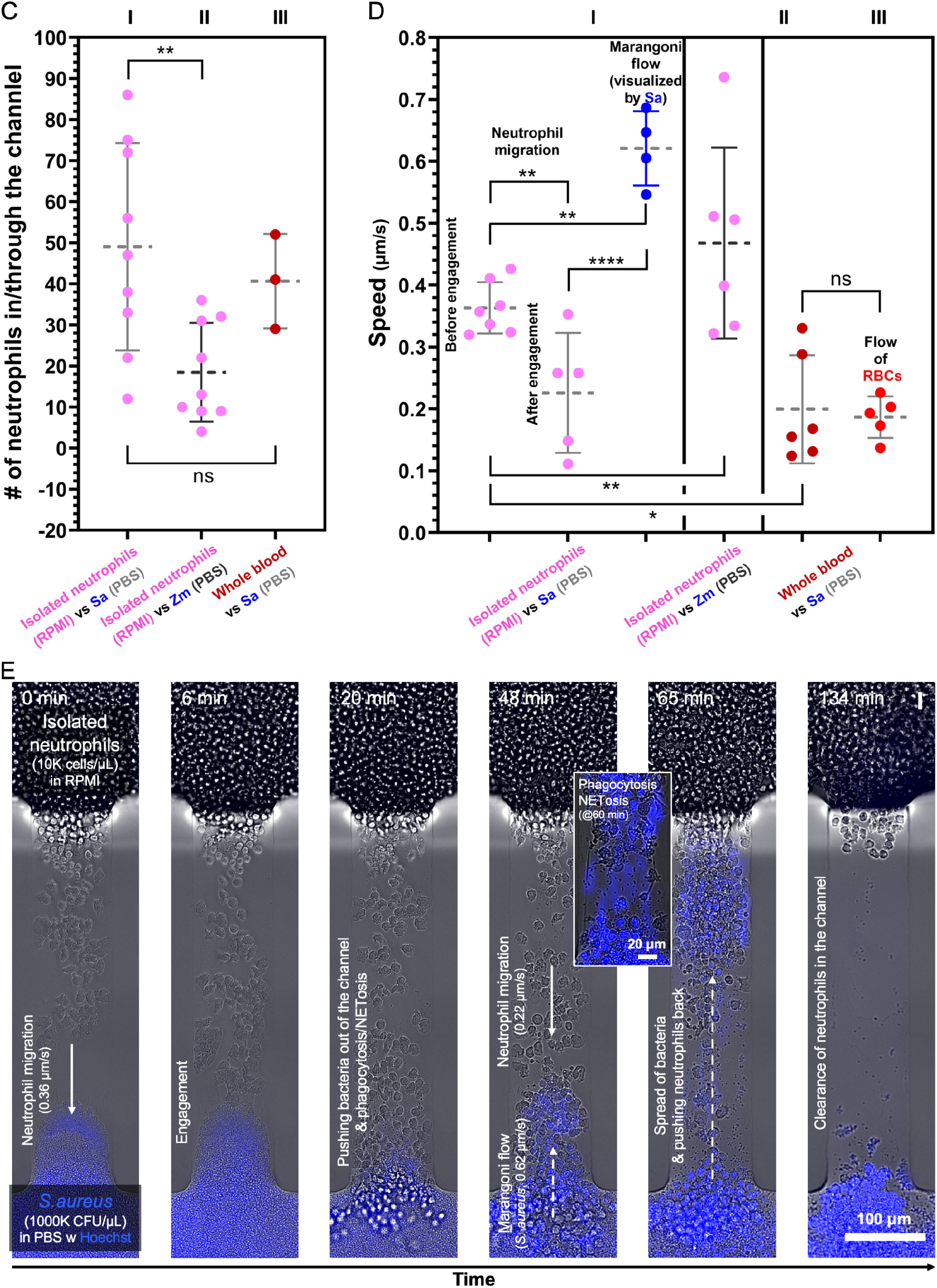
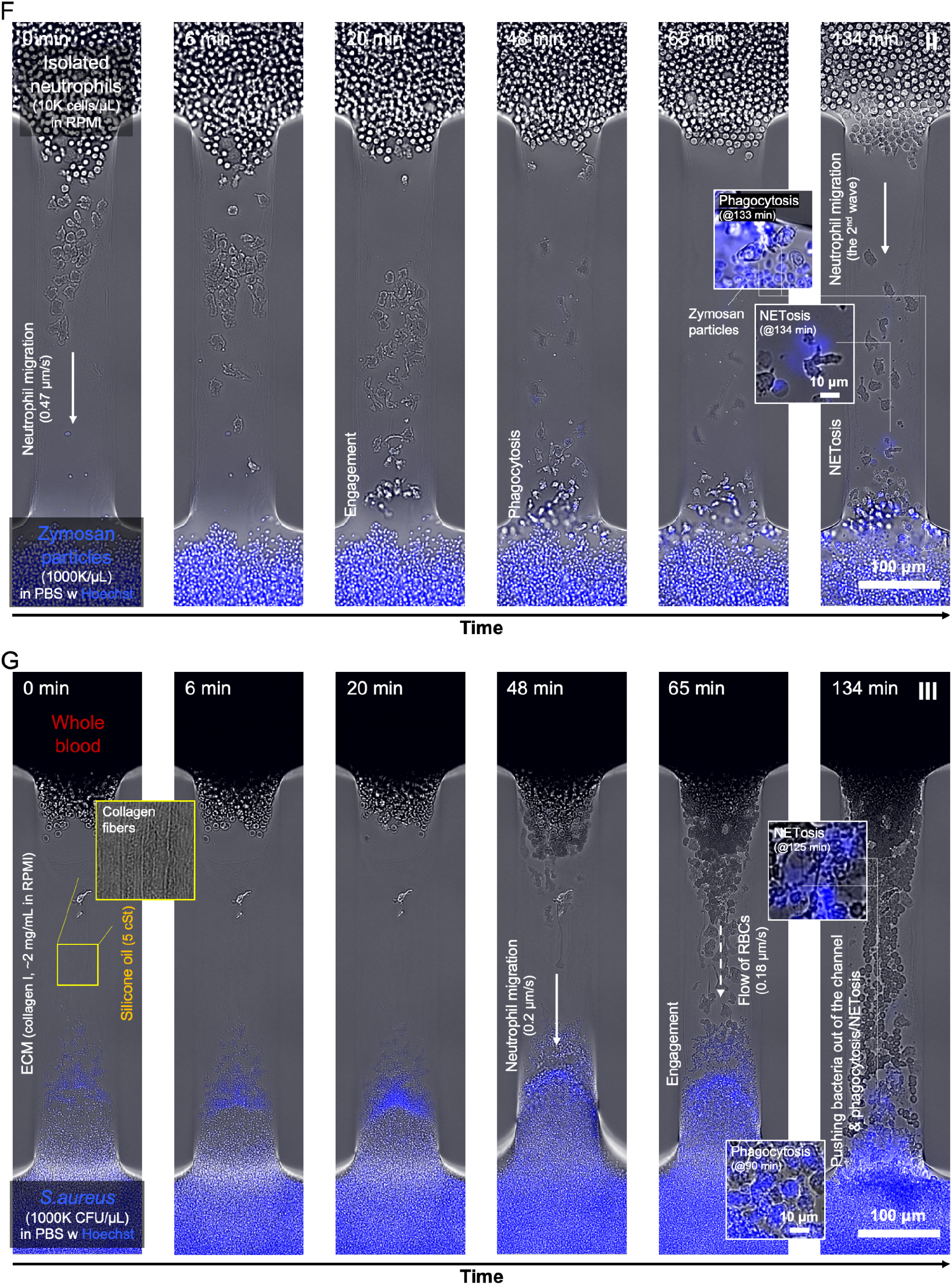
Generation of Marangoni flow with living bacteria versus dead yeast cells and isolated neutrophils versus whole blood. (A) A schematic showing the layout of the under-oil microchannels 2-2-*L*0.5-*W*0.1 (w ECM) and the initial interfacial tension of each liquid (Table S1). The immune niches (top) are isolated neutrophils (10K cells in 1 μL RPMI) or unprocessed whole blood (1 μL). The infection niches (bottom) are *S. aureus* (1000K CFUs in 1 μL PBS) or Zymosan particles (1000K in 1 μL PBS) with Hoechst (a DNA stain, BFP, 1:200 dilution). (B) Evolution of the estimated interfacial tension (Table S1) over time. The decrease rate of interfacial tension at the infection niche with *S. aureus* in PBS was adopted from literature for 3 mN/m/h.^[37]^ (C) The number of neutrophils in and through the channel and (D) speed (neutrophil migration, Marangoni flow, and flow of RBCs) results. (E), (F), and (G) The representative microscopic images (composite of bright-field and BFP) over 134 min of each condition in (A). (Insets) Close-ups of phagocytosis and NETosis visualized by Hoechst. Error bars are mean ± S.D. from ≥3 replicates. \**P* ≤ 0.05, \*\**P* ≤ 0.01, \*\*\**P* ≤ 0.001, and \*\*\*\**P* ≤ 0.0001. “ns” represents “not significant”.

In this experiment, we introduced a DNA stain (Hoechst) at the infection niche to visualize NETosis - the release of DNA content to the extracellular space by activated neutrophils (Movie S5). When neutrophils reached the infection niche, the generation and release of NETs were readily identified by the Hoechst stain (Figure 4E-G, insets). During migration in the microchannel, little to no NETosis was detected. The migrating neutrophils phagocytize the bacteria in the confined space and physically push the bacterial cells out of the space and back to the infection niche (Movie S2, Movie S5). These pathogen-control abilities identified in the single-cell confined space *in vitro* including phagocytosis, degranulation, purge of pathogen particles by pushback, and NETosis recapitulate the host defense enabled by neutrophils in response to invading bacteria. However, the successful pathogen control can get impaired and abolished in a few hours by Marangoni flow from the infection niche triggered by the growth of bacteria and secretion of biosurfactants.

### 2.5. Neutrophil-Bacteria Interaction in Confined Whole Blood Microenvironments

In standard *in vitro* immune-pathogen assays, the target immune cells are typically isolated from whole blood and interrogated in an artificial media. Immune cells are known with their inherently high sensitivity to the altered environmental factors [including nutrients, vital gas,^[40]^ autologous signaling molecules,^[41,42]^ and constituent cells^[43–46]^] from *in vivo* to *ex vivo*. Particularly, neutrophils get randomly and non-specifically activated (i.e., without operator-defined stimuli) simply by being pulled out of the autologous whole blood.^[22]^ Similarly, microorganisms have been reported showing significantly different growth kinetics and virulence when cultivated in different culture media.^[47–49]^ The development of the μ-Blood assay aims to perform immune cell functional assays and immune-pathogen assays with the target immune cells and microorganisms preserved and interrogated in the original whole blood microenvironment for improved extraction of donor-specific information and assay consistency.^[23]^

Here we test neutrophil response and the generation of Marangoni flow directly from a microliter of unprocessed whole blood in response to *S. aureus* in PBS (Figure 4C, D, and G), standard bacterial culture broth (MHB), and human serum (Figure 5). To get a complete whole blood environment in the microchannel, we adopted the microchannels 2-2-*L*0.5-*W*0.1 (w/o ECM) with the ECM layer removed considering that collagen I is only available from animal sources, e.g., murine, bovine. The microchannels without ECM coating require double-oil overlay (i.e., silicone oil + fluorinated oil) (Movie S6) to keep the microchannel hydrated with minimized water loss via evaporation under oil.^[23]^ In comparison, the microchannels with ECM coating are able to stay hydrated with single-oil overlay (i.e., silicone oil) (Figure 1).

**Figure 5.**
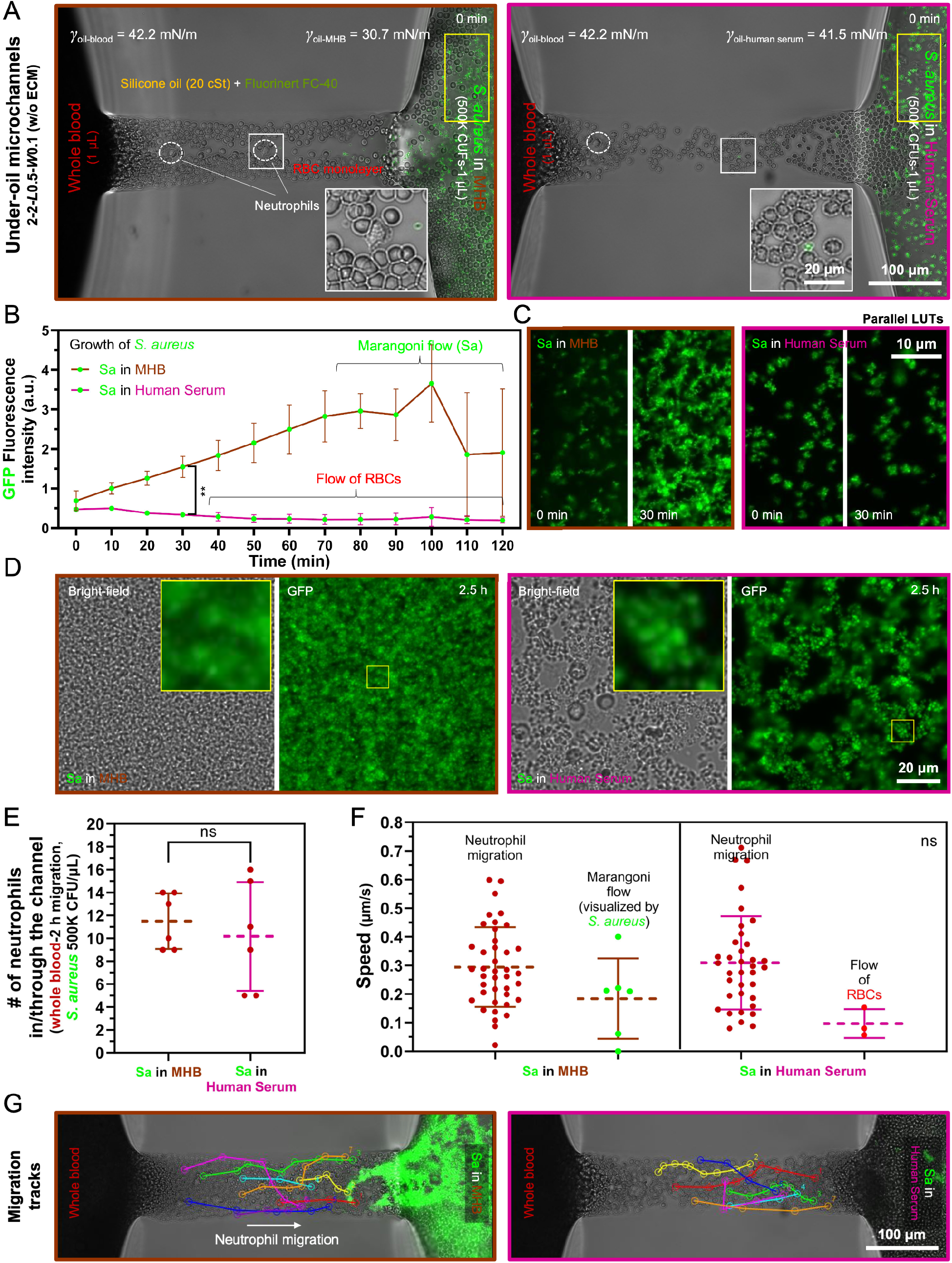
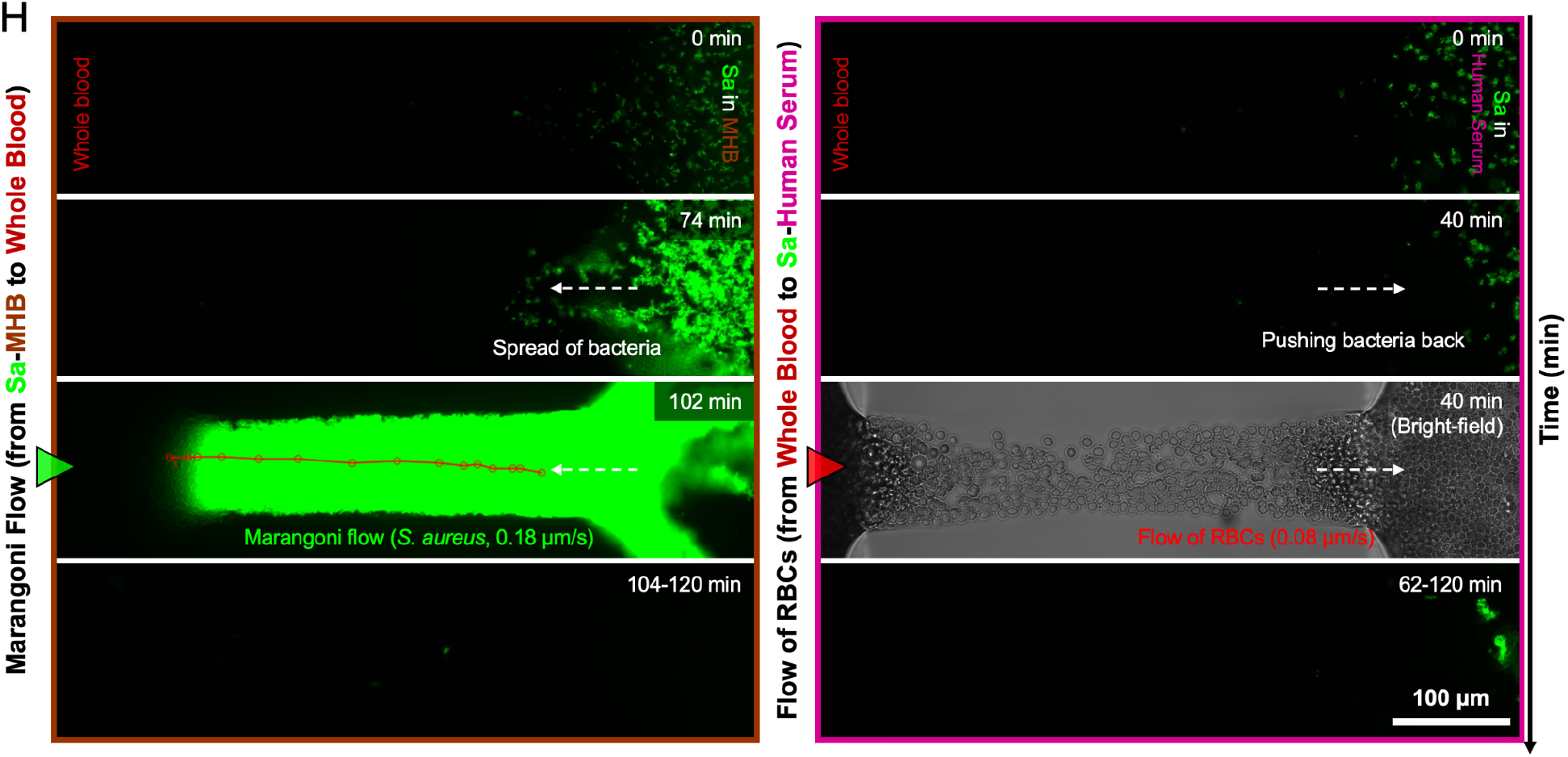
Neutrophil-bacteria interaction in confined whole blood microenvironments. (A) A top view (xy plane) of the under-oil microchannel [2-2-*L*0.5-*W*0.1 (w/o ECM) with the immune niche (1 μL whole blood) on the left and the infection niche (*S. aureus* in 1 μL MHB or human serum, 500K CFU/1 μL) on the right. (Insets) Zoom-in microscopic images showing the morphology of neutrophils against the RBC monolayer in the microchannel. (B) Growth of *S. aureus* at the infection niche spot [ROIs highlighted by the yellow solid-line boxes in (A)] monitored and quantified by GFP fluorescence intensity. The decrease of fluorescence intensity in the MHB group is caused by Marangoni flow, pumping the bacterial mass through the microchannel into the immune niche spot. In comparison, the decrease of fluorescence intensity in the human serum group is attributed to the flow of RBCs, pushing the bacterial mass backward at the infection niche spot. (C) and (D) The representative microscopic images showing the structure and different growth rates of *S. aureus* in MHB and human serum at 30 min and 2.5 h. (E) Neutrophil recruitment and (F) Speed analysis of neutrophil migration, Marangoni flow, and flow the RBCs. (G) The representative microscopic images showing the cell tracks. (H) The representative microscopic images showing Marangoni flow in the MHB group and the flow of RBCs in the human serum group. The Error bars are mean ± S.D. from ≥3 replicates. \**P* ≤ 0.05, \*\**P* ≤ 0.01, \*\*\**P* ≤ 0.001, and \*\*\*\**P* ≤ 0.0001. “ns” represents “not significant”.

First in response to *S. aureus* in PBS (*γ*_oil-PBS_ = 40.2 mN/m) (Table S1) in 2-2-*L*0.5-*W*0.1 (w ECM and single-oil overlay - silicone oil 5 cSt), neutrophils responded and migrated toward bacteria directly from whole blood (*γ*_oil-blood_ = 33.9 mN/m) at an average speed of 0.2 μm/s (Figure 4C, D, and G, Movie S5). In 134 min, no backflow was detected from the infection niche to the immune niche. Neutrophils pushed the bacteria spreading into the microchannel backward to the infection niche. The absence of backflow is attributed to the significantly lower interfacial tensions of whole blood compared to PBS (Figure 4A, B). It is worth noting that in this whole blood-versus-PBS condition Marangoni flow should exist pointing from the immune niche to the infection niche. However, due to the single-cell confinement with a channel height of 1.6 μm in the microchannel 2-2-*L*0.5-*W*0.1 (w ECM) (Figure 1C), the Marangoni flow could not be readily visualized. When neutrophils are migrating into the channel, the microchannel gets pushed open in z direction due to the less deformable nuclei of neutrophils (Figure 1C).^[50]^ The details were studied and reported in our pilot μ-Blood project.^[23]^ With the enlarged channel height, RBCs got carried over along with neutrophil migration independent from the Marangoni flow, forming a “flow of RBCs” that can be readily identified with the same direction of neutrophil migration and slower speed (Figure 4D and G, Movie S5).

Next we inoculated *S. aureus* in MHB (*γ*_oil-MHB_ = 30.7 mN/m) compared to human serum (*γ*_oil-human serum_ = 41.5 mN/m) (Table S1) in 2-2-*L*0.5-*W*0.1 (w ECM and double-oil overlay - silicone oil 20 cSt + Fluorinert FC-40) in response to whole blood (*γ*_oil-blood_ = 42.2 mN/m) (Figure 5A). Due to the increased media layer thickness (1.6 μm with ECM versus 2.3 μm without ECM) (Figure 1C), a monolayer of RBCs passively spreaded into the microchannel from whole blood loading on the immune niche spot. After sample loading, we run a 2-h timelapse (Movie S7). The first noticeable difference between MHB and human serum is the growth rate of *S. aureus* (Figure 5B-D). In MHB, *S. aureus* showed a quick growth. By contrast, *S. aureus* showed a much slower growth in human serum. In addition, in MHB *S. aureus* showed Brownian motion in a dense layer of bacteria but in human serum it formed a completely stationary biofilm-like network (Movie S8). The different growth rate of *S. aureus* in the two conditions didn’t cause a noticeable difference in neutrophil recruitment and migration speed (Figure 5E-G).

The other stark difference identified in this experiment between MHB and human serum is the backflow (Figure 5H, Movie S7). In the whole blood-versus-MHB condition, the quick growth of *S. aureus* and the low interfacial tension of the broth media (Figure 5A) triggered a strong backflow (Marangoni flow) starting at 74 min with a speed of 0.18 μm/s and peaking at a speed of 23.1 μm/s at 2.5 h (Movie S9), pumping the entire chunk of bacterial mass next to the entrance plume at the infection niche directly into the immune niche. In comparison, no backflow was detected in the whole blood-versus-human serum condition due to both the slow growth of *S. aureus* and the close interfacial tension between whole blood and human serum (Figure 5A). Instead, a flow of RBCs was formed - relatively slow in the beginning for 0.08 μm/s and then peaking at a speed of 4.7 μm/s at 2.5 h (Movie S9) - along with neutrophil migration, pushing the bacteria further back at the infection niche (Figure 5H, Movie S7).

Directly from a microliter of unprocessed whole blood enabled by μ-Blood, neutrophils are recruited toward invading bacteria in a space with single-cell confinement, showing a successful pathogen control at the initial stage (Figure 4G, Figure 5G). However, the final outcome of the war between neutrophils and bacteria in such a confined space is dominated unexpectedly by the spontaneously generated Marangoni flow at the infection niche. The invading bacteria gain a dominant advantage from the backflow when triggered, reversing the innate immune intervention and getting pumped directly into the immune niche.

## 3. Conclusion

*In vitro* immune-pathogen assays allow the interrogation of primary, patient-specific/matched samples (e.g., blood cells, microbial isolates) with improved physical, optical, and biochemical access compared to *in vivo* animal models. Multi-parametric readouts from molecular level to phenotypic level can be continuously extracted and analyzed, providing valuable information for diagnosis and treatment in personalized medicine. One long-existing challenge in *in vitro* immune-pathogen assays is the discrepancy between the *in vitro* assay environment and the *in vivo* physiopathological environment. The altered environmental factors cause random artifacts on both the target immune cells and microorganisms, tarnishing the extraction of donor-relevant/specific information and assay consistency.

The ELR-empowered UOMS leads a new sub-branch in open microfluidics with improved functionality including micrometer-scale lateral resolution (compared to millimeter-scale lateral resolution in other existing open microfluidic systems), open-fluid single-cell trapping, high flow rate range (comparable to blood flow in the circulatory systems), on-demand reversible open-fluid valves,^[51]^ and new functions [e.g., autonomously regulated oxygen microenvironment (AROM)].^[52]^ These advances in ELR-empowered UOMS together not only make open microfluidics comparable to the closed-chamber/channel microfluidics but also introduces new avenues to cellular^[24,25,51–54]^ and molecular^[55]^ assays in biology and biomedicine.

Enabled by the unique physics and functionality of ELR-empowered UOMS, we developed μ-Blood - an *in vitro* immune cell functional assay model that supports multiple phenotypic readouts of neutrophil function (including cell/nucleus morphology, motility, recruitment, and pathogen control) using a microliter of unprocessed whole blood. In this work, μ-Blood is extended to study and compare neutrophil-bacteria interactions in different exogenous and endogenous fluid microenvironments with single-cell confinement.

It is worth noting that in the standard assay with isolated neutrophils in RPMI versus bacteria in PBS the seeding density of neutrophils (10K cells/μL) has to be much higher than the physiological abundance of neutrophils in whole blood (with a healthy adult ranging from 2.5K to 7K cells/μL). As systematically studied and reported in our pilot μ-Blood project,^[23]^ the seeding density of neutrophils at the physiological level is limited by the random non-specific activation of neutrophils in RPMI with either animal or human serum supplements, leading to 10-70% non-specific activation in only 3 h and diminished (>90%) neutrophil recruitment compared to in whole blood for <0.1% non-specific activation and sustained neutrophil recruitment for more than 3 days. While using an artificially increased seeding density (10K cells/μL or above) is able to get neutrophil recruitment, it fails to recapitulate the physiologically relevant, donor-specific neutrophil abundance and kinetics. Similarly, microorganisms are known to utilize fast mutation to adapt themselves to the changing environment. Different culture media or blood/serum from different donors significantly alter the growth kinetics and virulence.^[47–49]^ Additionally, the typical *in vivo* cellular microenvironments (e.g., capillary bed) features highly heterogeneous, dynamic, and confined structures and mass transport at single-cell level and micrometer scale. The *in vivo* cellular microenvironments are fundamentally different compared to the bulk-scale assays that are typically performed in an open space (e.g., Petri dish, microtiter plate, glass slide/coverglass) with minimal single-cell level confinement. These insights highlight the necessity of performing *in vitro* immune-pathogen assay directly in an endogenous fluid microenvironment (e.g., whole blood) with single-cell confinement.

In the confined whole blood microenvironments, neutrophil intervention in response to the invading bacteria is predominantly influenced by the capillary convection known as Marangoni flow. The trigger of Marangoni flow at the infection niche is determined by the growth kinetics of bacteria and the secretion of biosurfactants. It needs to be noted that the generation and tracking of Marangoni flow in μ-Blood are visualized by the microparticles (including bacteria, neutrophil granule-bacteria complex, ECM fibers) getting carried in the flow. The starting time of Marangoni flow should be earlier than the time of the flow being detectable with the microparticles. The exact fluid dynamics of generation and evolution of Marangoni flow can be achieved by numerically simulating the under-oil microchannels.

In the future work aiming for translational applications, μ-Blood can used to screen whole blood samples from a donor pool including healthy, cancer, autoimmune, and infection diseases, in Phase I - in response to a library of laboratory human pathogens including bacterial species that cause bacteremia, septicemia, and pneumonia (e.g., *Pseudomonas aeruginosa*, *Staphylococcus aureus*, *Klebsiella pneumoniae*, *Acinetobacter baumannii*, *Enterobacter species*, *Extra-intestinal pathogenic Escherichia coli*, *Streptococcus agalactiae*, *Streptococcus pneumoniae*, and *Haemophilus influenzae*), fungal species that cause systemic fungal infection diseases (e.g., *Candida albicans*, *Aspergillus fumigatus*), and fungal-bacterial mix that used to result in the worst treatment outcomes; in Phase II - in response to clinical microbial isolates with whole blood from the matched patient with different antimicrobial intervention strategies (e.g., combination antibiotic therapy, sequential therapy). A dominant advantage of μ-Blood compared to other microfluidics-based immune-pathogen assay platforms is its open microfluidic configuration. μ-Blood allows free physical access to the samples on the device with minimized system disturbance (including evaporation and contamination) and naturally aligns with automated liquid handling systems.^[56]^ Flexible and high-throughput spatiotemporal sample manipulation and collection for large-sample-size screening and further off-chip, downstream analyses can be readily performed with μ-Blood, e.g., RNA sequencing for studying inherent immune cell heterogeneity and discovery of new resistant genes in pathogens.

## 4. Experimental Section

### Preparation of ELR-empowered UOMS devices

<u>Step #1)</u> Fabrication of PDMS silane-grafted surface. Chambered coverglass [Nunc Lab-Tek-II, 2 well (155379), #1.5 borosilicate glass bottom, 0.13 to 0.17 mm thick; Thermo Fisher Scientific] (Figure 1b) was treated first with O_2_ plasma (Diener Electronic Femto, Plasma Surface Technology) at 100 W for 3 min and then moved to a vacuum desiccator (Bel-Art F420220000, Thermo Fisher Scientific, 08-594-16B) for chemical vapor deposition (CVD). PDMS-silane (1,3-dichlorotetramethylsiloxane; Gelest, SID3372.0) (25 μL ×2 per treatment) was vaporized under pumping for 3 min and then condensed onto glass substrate under vacuum at room temperature for 40 min. The PDMS-grafted surface was thoroughly rinsed with ethanol (anhydrous, 99.5%), deionized (DI) water, and then dried with nitrogen for use. <u>Step #2)</u> Fabrication of PDMS stamps. Photomasks were designed in Adobe Illustrator and then sent to a service (Fineline Imaging) for printing. Standard photolithography was applied to make a master that contains all the microchannel features. Details about photolithography can be found in our previous publication^[26]^. Last, PDMS stamps were made by pouring a degassed (20 min using a vacuum desiccator) silicone precursor and curing agent mix (SYLGARD 184, Silicone Elastomer Kit, Dow, 04019862) in 10:1 mass ratio onto the master and cured on a hotplate at 80 °C for 4 h. The PDMS stamps were stripped off with tweezers and punched with through holes (Miltex Biopsy Punch with Plunger, Ted Pella, 15110) at the inlet and outlet of a microchannel for the following O_2_ plasma surface patterning. <u>Step #3)</u> O_2_ plasma surface patterning. The PDMS silane-grafted chamber coverglass was masked by a punched PDMS stamp and then treated with O_2_ plasma at 100 W for 1 min. After surface patterning, the PDMS stamp was removed by tweezers and stored in a clean container (e.g., Petri dish) for reuse. <u>Step #4-1)</u> Under-oil microchannels without ECM coating - The chemically patterned chambered coverglass from Step #3 was overlaid with 1 mL silicone oil (<100 cSt, e.g., 5 cSt or 20 cSt) for each well on the 2-well chambered coverglass. The target media was distributed onto the microchannels by under-oil sweep (Figure 1a). Briefly, get 20 μL of the target media in a 1-200 μL large orifice pipette tip (02-707-134, Thermo Fisher Scientific) and then drag the hanging microdrop at the end of the tip through the patterned surface. Media was spontaneously distributed onto the O_2_ plasma-treated areas only. For double-oil overlay [i.e., silicone oil (20 cSt) + fluorinated oil (Fluorinert FC-40)], we first prepared the under-oil microchannels under silicone oil (20 cSt) with the target media by under-oil sweep. Next, we added 1 mL Fluorinert FC-40 (1.85 g/mL at 25 °C) for each well on the 2-well chambered coverglass directly into the silicone oil (20 cSt, 0.95 g/mL at 25 °C). Due to the high density of the fluorinated oil and immiscibility with silicone oil, the fluorinated oil spontaneously replaces silicone oil in a well and pushes it to the top layer. At last, the silicone oil was removed using pipet at the four corners in a well. <u>Step #4-2)</u> Under-oil microchannels with ECM coating - Similar to Step#4, the under-oil microchannels with ECM coating were prepared by sweeping a hanging microdrop of the ECM solution under oil (silicone oil, 5 cSt). The ECM solution is collagen I (Rat Tail, 10 mg/mL in the original bottle) 1:1 dilution in 2% N-2-hydroxyethylpiperazine-N-2-ethane sulfonic acid (HEPES) 2× PBS and then 1:1 dilution in RPMI (basal media only). The final pH of the collagen solution was adjusted to 7.4 by 0.5 M sodium hydroxide (NaOH) endotoxin-free aqueous solution (0.5 μL of 0.5 M NaOH per 100 μL collagen solution). After under-oil sweep, the device was kept at room temperature (∼21 °C) for 1 h for polymerization. The ECM layer effectively retains hydration of the microchannels with single-oil overlay. After the ECM layer is fully polymerized, the device is ready for sample loading. <u>Step #5)</u> Sample loading before imaging. The device with oil overlay was set up under a microscope. We first registered the xy and z positions of the microchannels with perfect focal plane (PFS) function. Next, we loaded the blood or cell stock using a pipet under oil for 1 μL/spot (Figure 1A) throughout all the microchannels. At last, we loaded the pathogen inoculum at the other end of each microchannel for 1 μL/spot. After sample loading and update of the xy and z positions, the device is ready for imaging or timelapse.

### Whole blood collection

All blood samples were drawn according to Institutional Review Boards (IRB) approved protocols per the Declaration of Helsinki at the University of Wisconsin-Madison in the Microtechnology, Medicine, and Biology (MMB) Lab (IRB# 2020-1623, healthy donors) and in the Lang Lab (IRB# 2014-1214, cancer patients). Informed consent was obtained from all subjects in the study. Whole blood was collected with standard Vacutainer tubes (ACD tube, BD, 364606). All the whole blood samples were stored in a standard CO_2_ incubator (Thermo Fisher Scientific, HERACELL VIOS 160i) at 37 °C before use.

### Neutrophil isolation

Neutrophils were isolated from whole blood using magnetic bead-based negative selection - EasySep Direct Human Neutrophil Isolation Kit (STEMCELL, 19666). RPMI (basal media only) was used as the isolation buffer. Cell count of each isolation was obtained using a hemocytometer (LW Scientific).

### Human serum separation

Human serum was separated from the same whole blood sample along with neutrophil isolation using the standard Vacutainer tubes (Serum Separation Tube, BD, 367989). The serum was aliquoted and stored at a −80 °C freezer (Thermo Fisher Scientific, Forma 900 Series) for future use. The aliquots needed in an experiment were thawed at room temperature (∼21 °C) with only one freeze-thaw cycle.

### Bacteria growth condition

Bacterial strains stored at −80 °C were plated on Mueller–Hinton agar (MHA, BD Difco, BD, Franklin Lakes, NJ USA) and incubated at 37 °C overnight prior to use. The inoculum was prepared to the starting bacterial concentration before an experiment. Zymosan (*S. cerevisiae*) BioParticles were purchased (Thermo Fisher Scientific, Z23373) and reconstituted in PBS prior to use.

### Microscopic imaging

Bright-field, fluorescence images and videos were recorded on a Nikon Ti Eclipse inverted epifluorescence microscope (Nikon Instruments). The chambered coverglass with oil overlay was kept at 37 °C, 21% O_2_, 5% CO_2_, and 95% RH in an onstage incubator (Bold Line, Okolab) during imaging or timelapse. After imaging on the microscope, the μ-Blood devices were moved to and kept in a standard CO_2_ incubator (37 °C, 18.6% O_2_, 5% CO_2_, and 95% RH) before the characterization for the next time point. Movies were prepared in the VSDC Video Editor.

### Cell counting and speed tracking analysis

Manual counting was performed in Fiji ImageJ with the timelapse videos recorded on the Nikon microscope by counting each individual neutrophil that passes into the channel. For speed tracking, images in a timelapse were transferred into Fiji ImageJ, where cell tracking was performed manually using the Fiji plugin “MTrackJ”. Once “MTrackJ” is selected, the “add” option allows tracking of each individual migrating cell. Once the manual tracking is done for each migrating cell, measurements can be taken automatically. Measurements were created through the “MTrackJ” measurements option.

### Statistical analysis

Raw data was directly used in statistical analysis with no data excluded. Data was averaged from at least 3 replicates (unless otherwise stated) and present as mean ± standard deviation (S.D.) if applicable. The statistical significance was specified in the figure captions. All statistical analyses were performed using GraphPad Prism 10.0.1.

## Acknowledgements

Experimental work was performed in the MMB Lab facility at the University of Wisconsin-Madison. We thank Dr. David J. Beebe for offering suggestions and Mrs. Sue McCrone (The Rose Lab at the University of Wisconsin-Madison) for providing the standard *S. aureus* inoculum. This work was supported by NIH P30 CA014520, NIH U24 AI152177, NIH R01 AI34749, and NH R01 AI54940. Author contributions: C.L. developed the under-oil open microfluidic system and designed the *in vitro* neutrophil-bacteria assay. C.L. performed data collection, analysis, and visualization with the assistance from N.W.H., Z.A., M.M., and J.-S.K.. C.L. supervised the project. C.L. wrote the manuscript and all authors revised it.

## Data Availability Statement

The data that support the findings of this study are available from the corresponding authors upon reasonable request.

