## Supporting Information for "*In Vitro* Neutrophil-Bacteria Assay in Whole Blood Microenvironments with Single-Cell Confinement"

**This PDF includes**

Table S1
Figures S1 and S2
Movies S1 to S9

| | Surface tension (air), mN/m | Interfacial tension* (silicone oil 5 cSt), mN/m | Interfacial tension* (Fluorinert FC-40), mN/m |
|---|---|---|---|
| Deionized (DI) water | 72.2 (25 °C)<br>69.5 (25 °C) (47) | 41.8 (21 °C) (24) | 52.1 (21 °C) (24) |
| Phosphate buffered saline (PBS) | 69.5 (25 °C) (47) | 40.2 (25 °C) | / |
| Basal medium | 69 (24 °C) (48) | 39.9 (24 °C) | / |
| Whole blood | 58.5 (22 °C)<br>52 (37 °C) (49) | 33.9 (22 °C) | 42.2 (22 °C) |
| Human serum | 57.5 (22 °C)<br>52 (37 °C) (49) | / | 41.5 (22 °C) |
| Mueller Hinton Broth (MHB) (beef infusion broth) | 46.2 (23 °C) (50) | / | 30.7 (23 °C) |

**Table S1. A summary of surface tension and interfacial tension of the liquids in this study.** *The interfacial tensions under oil are estimated proportionally based on the DI water values. For example, interfacial tension of silicone oil-PBS = (interfacial tension of silicone oil-water/surface tension of water) × surface tension of PBS.

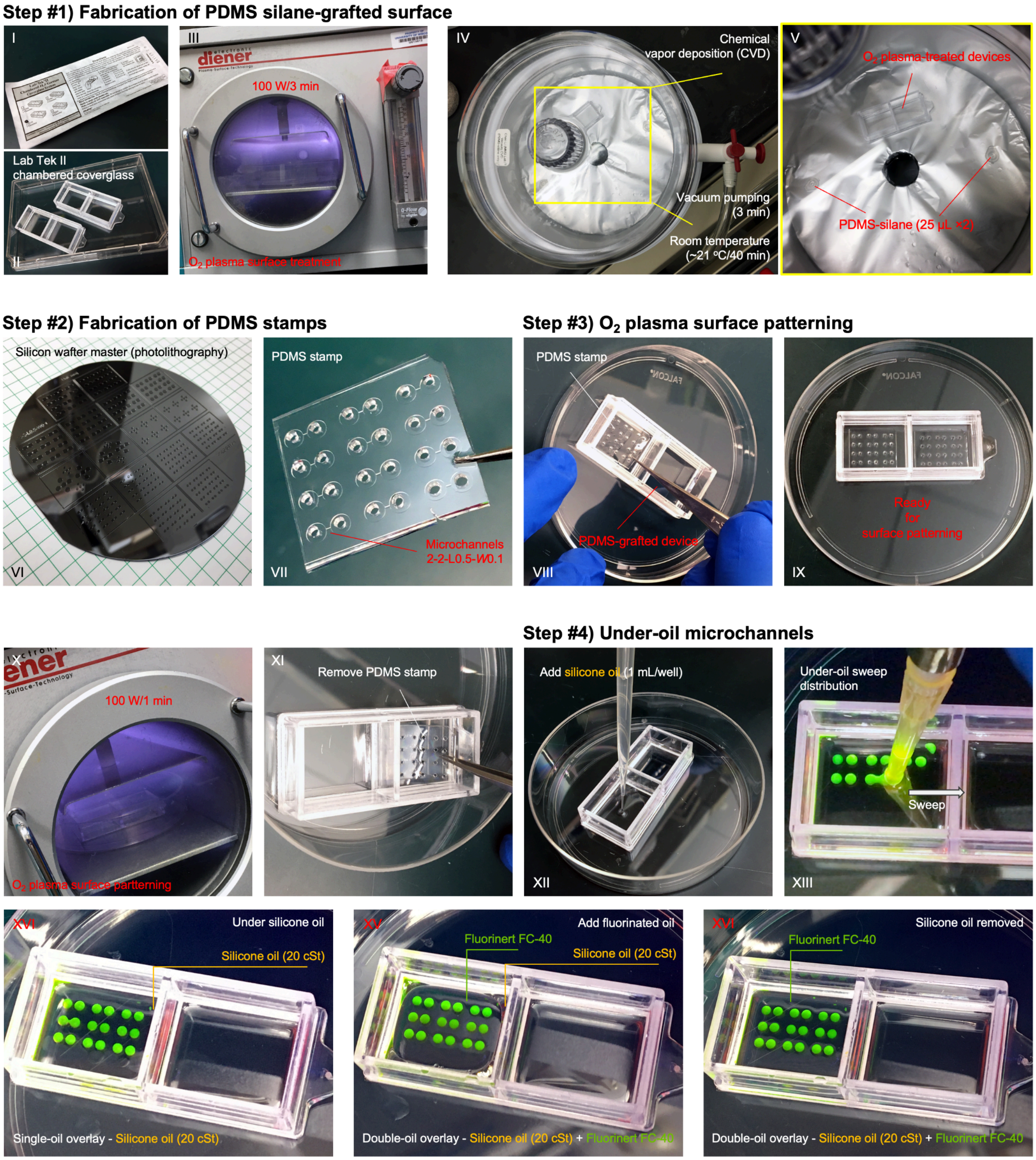
Step #1) Fabrication of PDMS silane-grafted surface
I
Lab Tek II
chambered coverglass
II
III
100 W/3 min
$O_2$ plasma surface treatment
IV
Chemical
vapor deposition (CVD)
Vacuum pumping
(3 min)
Room temperature
(~21 °C/40 min)
V
$O_2$ plasma-treated devices
PDMS-silane (25 µL ×2)
Step #2) Fabrication of PDMS stamps
Silicon wafter master (photolithography)
VI
PDMS stamp
Microchannels
2-2-L0.5-W0.1
VII
Step #3) $O_2$ plasma surface patterning
PDMS stamp
PDMS-grafted device
VIII
Ready
for
surface patterning
IX
X
100 W/1 min
$O_2$ plasma surface partterning
XI
Remove PDMS stamp
Step #4) Under-oil microchannels
Add silicone oil (1 mL/well)
XII
Under-oil sweep
distribution
Sweep
XIII
XVI
Under silicone oil
Silicone oil (20 cSt)
Single-oil overlay - Silicone oil (20 cSt)
XV
Add fluorinated oil
Fluorinert FC-40
Silicone oil (20 cSt)
Double-oil overlay - Silicone oil (20 cSt) + Fluorinert FC-40
XVI
Silicone oil removed
Fluorinert FC-40
Double-oil overlay - Silicone oil (20 cSt) + Fluorinert FC-40

**Step #5) Sample loading & imaging/timelapse**

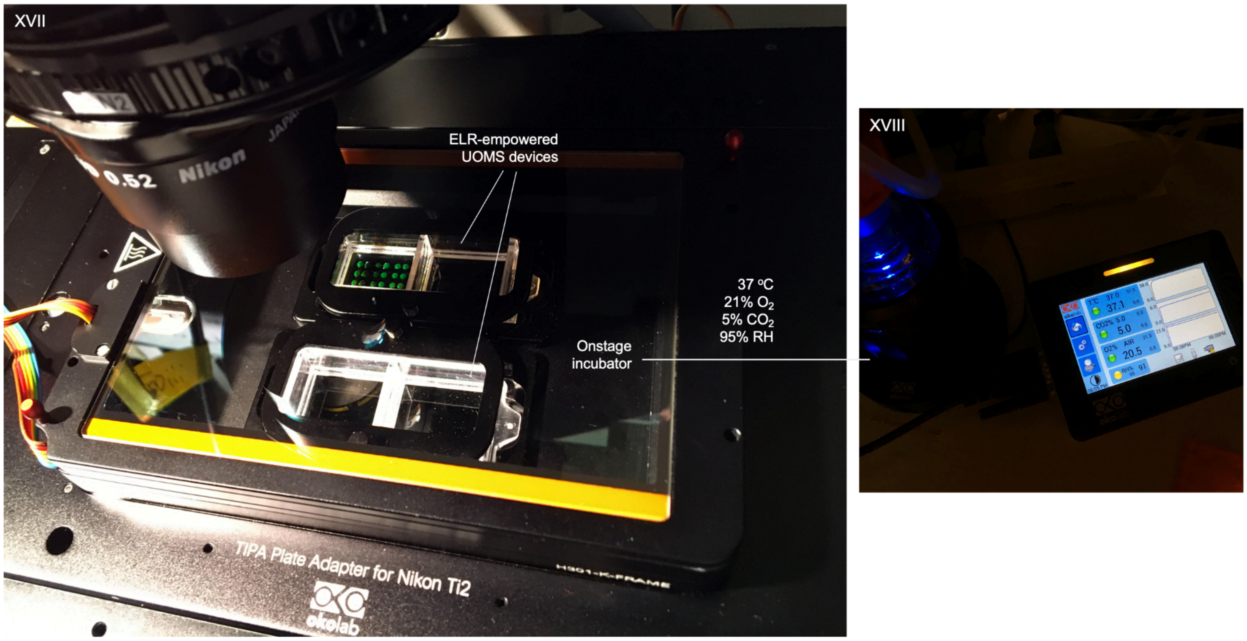


**Figure S1.** Workflow of ELR-empowered UOMS device fabrication and imaging setup. See detailed description of the steps in the Experimental Section.

A **Chambered coverglass**

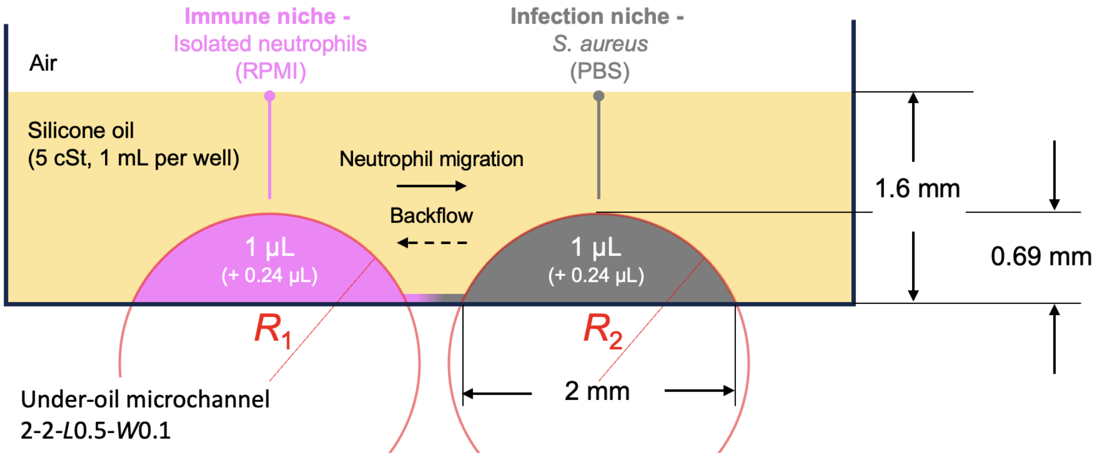


B **If the backflow is Laplace pressure-driven flow** **(i.e., high interfacial tension to low interfacial tension)**

Laplace pressure $\Delta P = 2\gamma_{\text{oil-media}}/R$

where $\gamma_{\text{oil-media}}$ is the interfacial tension at the oil-media interface, $R$ is the radius of curvature

($\gamma_{\text{oil-RPMI}}$ = 39.9 mN/m, $\gamma_{\text{oil-PBS}}$ = 40.2 mN/m, $R_1 = R_2$ = 1.07 mm)

**then $\gamma_{\text{oil-RPMI}}$ needs to decrease and/or $\gamma_{\text{oil-PBS}}$ needs to increase over time.**

C **If the backflow is Marangoni flow** **(i.e., low interfacial tension to high interfacial tension)**

**then $\gamma_{\text{oil-RPMI}}$ needs to increase and/or $\gamma_{\text{oil-PBS}}$ needs to decrease over time.**

**Figure S2.** Mechanism analysis of the backflow observed in Figure 2. (A) A Schematic showing the under-oil fluid environment and relevant parameters. (B) Hypothesis of Laplace pressure-driven flow with the flow direction from high interfacial tension to low interfacial tension. Calculation of the Laplace pressure in the sessile microdrop on the two spots is in the brackets. (C) Hypothesis of Marangoni flow with the flow direction from low interfacial tension to high interfacial tension.

**Movie S1.** Double-ELR and under-oil sweep distribution in ELR-empowered UOMS.

**Movie S2.** Neutrophil response and pathogen-control events in the standard assay fluid microenvironment - isolated neutrophils in RPMI versus *S. aureus* in PBS - in 2 h from sample loading.

**Movie S3.** Comparison of Marangoni flow between *S. aureus* 1000K CFU/μL and 100K CFU/μL inoculum density at 160 min from sample loading.

**Movie S4.** Real-time neutrophil migration in response to *S. aureus* 25K CFU/μL inoculum density at 140 min from sample loading.

**Movie S5.** Comparison between isolated neutrophils (in RPMI) and whole blood in response to live bacteria and dead yeast cell particles (in PBS).

**Movie S6.** Double-oil overlay and operation in ELR-empowered UOMS.

**Movie S7.** Neutrophil response and pathogen-control events in the whole blood microenvironments - neutrophils in whole blood versus *S. aureus* in MHB and human serum - in 2 h from sample loading.

**Movie S8.** Comparison of *S. aureus* motion at 2.5 h after sample loading in MHB and human serum.

**Movie S9.** Marangoni and RBC flow in the whole blood microenvironments at 2.5 h from sample loading.
